# PTEN-induced kinase 1 gene single-nucleotide variants as biomarkers in adjuvant chemotherapy for colorectal cancer

**DOI:** 10.1101/2022.11.28.518259

**Authors:** Yoshiaki Mihara, Masataka Hirasaki, Yosuke Horita, Takashi Fujino, Hisayo Fukushima, Yasuo Kamakura, Kousuke Uranishi, Yasumitsu Hirano, Shomei Ryozawa, Masanori Yasuda, Yoshinori Makino, Satomi Shibasaki, Tetsuya Hamaguchi

## Abstract

**Background:** Fluoropyrimidine-based adjuvant chemotherapy is globally recommended for postoperative stage III colon cancer and high-risk stage II patients. However, adjuvant chemotherapy is often associated with severe adverse events and is not highly effective in preventing recurrence. Therefore, a recurrence-prevention biomarker of adjuvant chemotherapy for colorectal cancer is necessary for providing such treatments to appropriate patients.

Autophagy (including mitophagy) is activated under chemotherapy-induced stress and contributes to chemotherapy resistance. Expression of autophagy-related genes and their single-nucleotide polymorphisms are reported to be effective predictors of chemotherapy response in some cancers.

Our goal was to evaluate the relationship between the single-nucleotide variants of autophagy-related genes and recurrence rates to identify the recurrence-prevention biomarkers of adjuvant chemotherapy in colorectal cancer.

**Methods:** We analyzed surgical or biopsy specimens from 84 patients who underwent radical surgery followed by fluoropyrimidine-based adjuvant chemotherapy at Saitama Medical University International Medical Center between January and December 2016. Using targeted enrichment sequencing, we identified single-nucleotide variants and insertions/deletions in 50 genes, including autophagy-related genes, and examined their association with colorectal cancer patient relapse rates.

**Results:** We detected 560 single-nucleotide variants or insertions/deletions in the target region. The results of Fisher’s exact test indicated that the recurrence rate of colorectal cancer after adjuvant chemotherapy was significantly lower in patients with the single-nucleotide variants (c.1018G>A [*p* < 0.005] or c.1562A>C [*p* < 0.01]) of the mitophagy-related gene PTEN-induced kinase 1.

**Conclusions:** The two single-nucleotide variants of this mitophagy-related gene may be biomarkers of non-recurrence in colorectal cancer patients who received postoperative adjuvant chemotherapy.

## INTRODUCTION

Colorectal cancer (CRC) is the second leading cause of cancer-related death worldwide.^1^ Considering that the recurrence rate of stage III and high-risk stage II CRC is more than 30%, a fluoropyrimidine (5-FU)-based regimen is recommended as postoperative adjuvant chemotherapy.^2, 3^ However, diagnosis of high-risk stage II CRC is challenging because the criteria vary depending on the guidelines of different societies, such as American Society of Clinical Oncology, European Society for Medical Oncology, and National Comprehensive Cancer Network.^4-6^ Circulating tumor DNA has been postulated as a prognostic factor for the risk of postoperative recurrence in stage II colon cancer but has not been put into practical use because of its cost and the lack of sufficient evidence (the study subjects were exclusively stage II, and the observation period was insufficient).^7, 8^ Furthermore, the effectiveness of current adjuvant chemotherapy is not satisfactory. 5-FU-based adjuvant chemotherapy without oxaliplatin reduces the recurrence rate by only 10% compared with surgery alone, with a relative risk reduction of approximately 17%–32%.^2, 9^ Even with the addition of oxaliplatin to adjuvant chemotherapy, the recurrence rate is reduced by only 5%.^3^ 5-FU-based chemotherapy has also proven to be problematic as it causes severe toxicity in up to 30% of its recipients.^10, 11^ Therefore, the decision to use adjuvant chemotherapy in CRC is left to the attending physician.^12^ Given the above information, finding a recurrence-prevention biomarker in adjuvant chemotherapy for CRC is necessary to expedite and guide treatment decisions.

*KRAS* and *BRAF* mutations have been reported as possible predictors of poor prognosis for adjuvant chemotherapy in stage II/III colon cancer by a systematic review of nine randomized controlled phase III trials. However, *KRAS* mutations significantly worsened both disease-free survival (DFS) and overall survival (OS), whereas *BRAF* mutations worsened OS but did not worsen DFS significantly. In addition, the trials did not include rectal cancer^13^; therefore, our goal is to find a novel recurrence biomarker in adjuvant chemotherapy for CRC.

Autophagy is a highly regulated process that degrades and recycles cellular components. The most important features of autophagy include the breakdown of proteins and organelles in the cell and recycling them as a new source of nutrition.^14^ In human colon cancer cell lines, autophagy activation is induced by 5-FU treatment, and the inhibition of autophagy significantly increases 5-FU-induced apoptosis. Therefore, autophagy is activated as a protective mechanism against 5-FU-induced apoptosis.^15^ Mitophagy is a form of autophagy that allows mitochondria to maintain homeostasis and plays a role in the late stages of tumorigenesis by increasing cell resistance and promoting carcinogenesis. This process has been reported to mediate chemotherapy resistance in various types of cancer.^16^ The silencing of the BCL2/adenovirus E1B 19- kDa protein-interacting protein 3 (*BNIP3*) in CRC and the high expression of PTEN-induced putative kinase 1 (*PINK1*) in esophageal cancer are associated with resistance to 5-FU-based chemotherapy.^17, 18^ Both *BNIP3* and *PINK1* are mitophagy-related genes. We planned to establish a system to predict the efficacy of postoperative 5-FU-based adjuvant chemotherapy in CRC using the single-nucleotide variants (SNVs) of autophagy- and mitophagy-related genes.

The aim of this study was to find new recurrence-prevention biomarkers by analyzing autophagy- and cancer-related genes in specimens from patients undergoing 5-FU-based adjuvant chemotherapy for CRC and examine the association between the results and recurrence.

## MATERIALS AND METHODS

### Tissue samples

A total of 84 analytic samples from surgical or biopsy specimens were collected from 84 patients who underwent radical surgery for CRC at Saitama Medical University International Medical Center (SMUIMC) between January and December 2016. One case was excluded because the specimen was too small; therefore, we used a metastatic lymph node instead of the primary tumor. In three cases, double carcinoma was observed. In these situations, the specimen with the greatest depth was selected or if the depths were the same, the one with a lower differentiation was selected. The tumor area was confirmed by pathologists using the naked eye and microscopy.

All patients underwent curative surgery followed by postoperative 5-FU-based adjuvant chemotherapy. Postoperative adjuvant chemotherapy consisted of regimens based on 5-FU: S-1; capecitabine; tegafur–uracil plus leucovorin calcium; or oxaliplatin combinations such as FOLFOX (5-FU, levofolinate, and oxaliplatin), CAPOX (capecitabine and oxaliplatin), and SOX (S-1 and oxaliplatin). Recurrence was defined in terms of the date when CRC recurrence was confirmed via imaging, endoscopy, or clinical examination. The follow-up period for the presence of recurrence was within 5 years after surgery. Clinical information was obtained by reviewing medical records and pathology reports (Table 1).

**TABLE 1.**
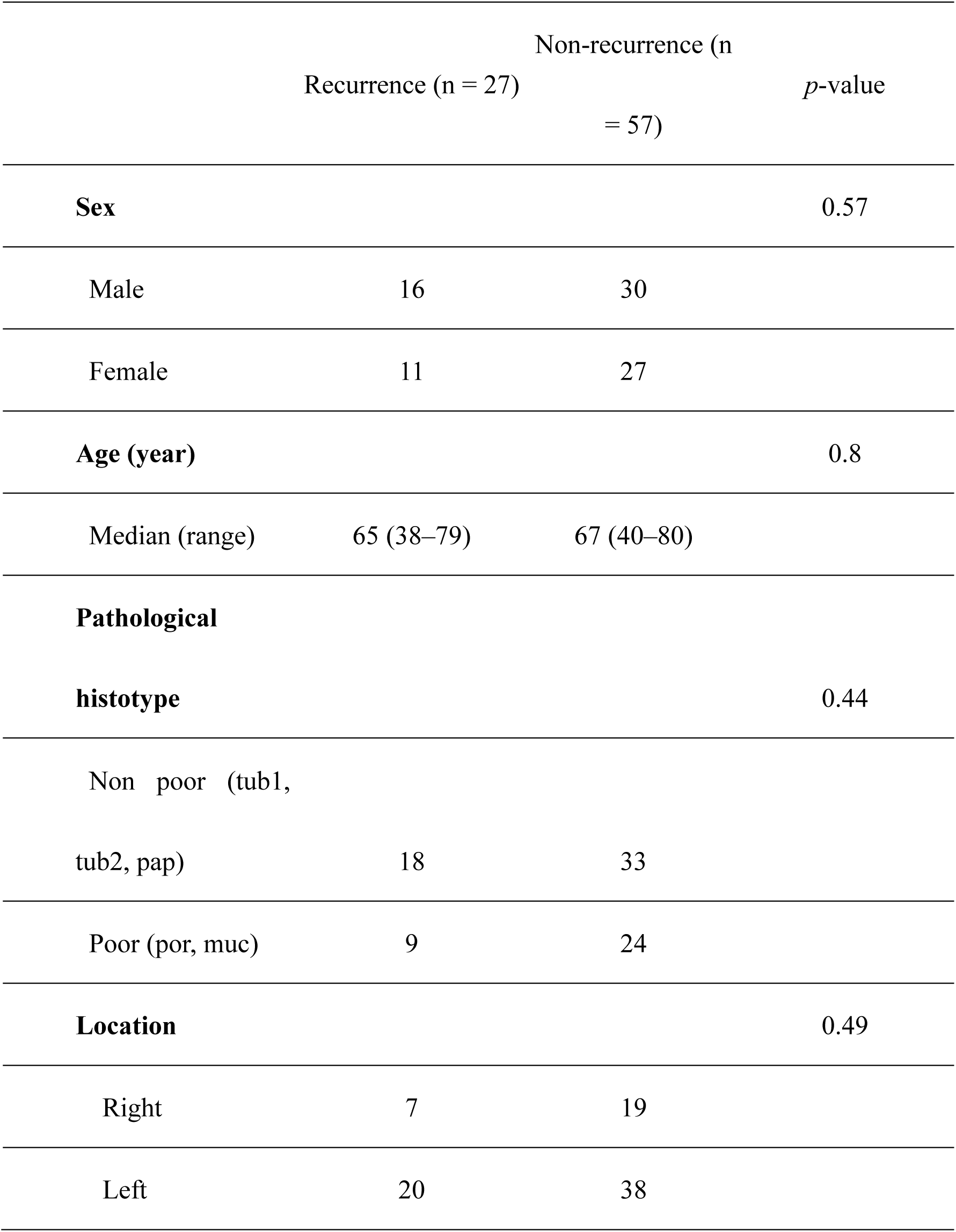
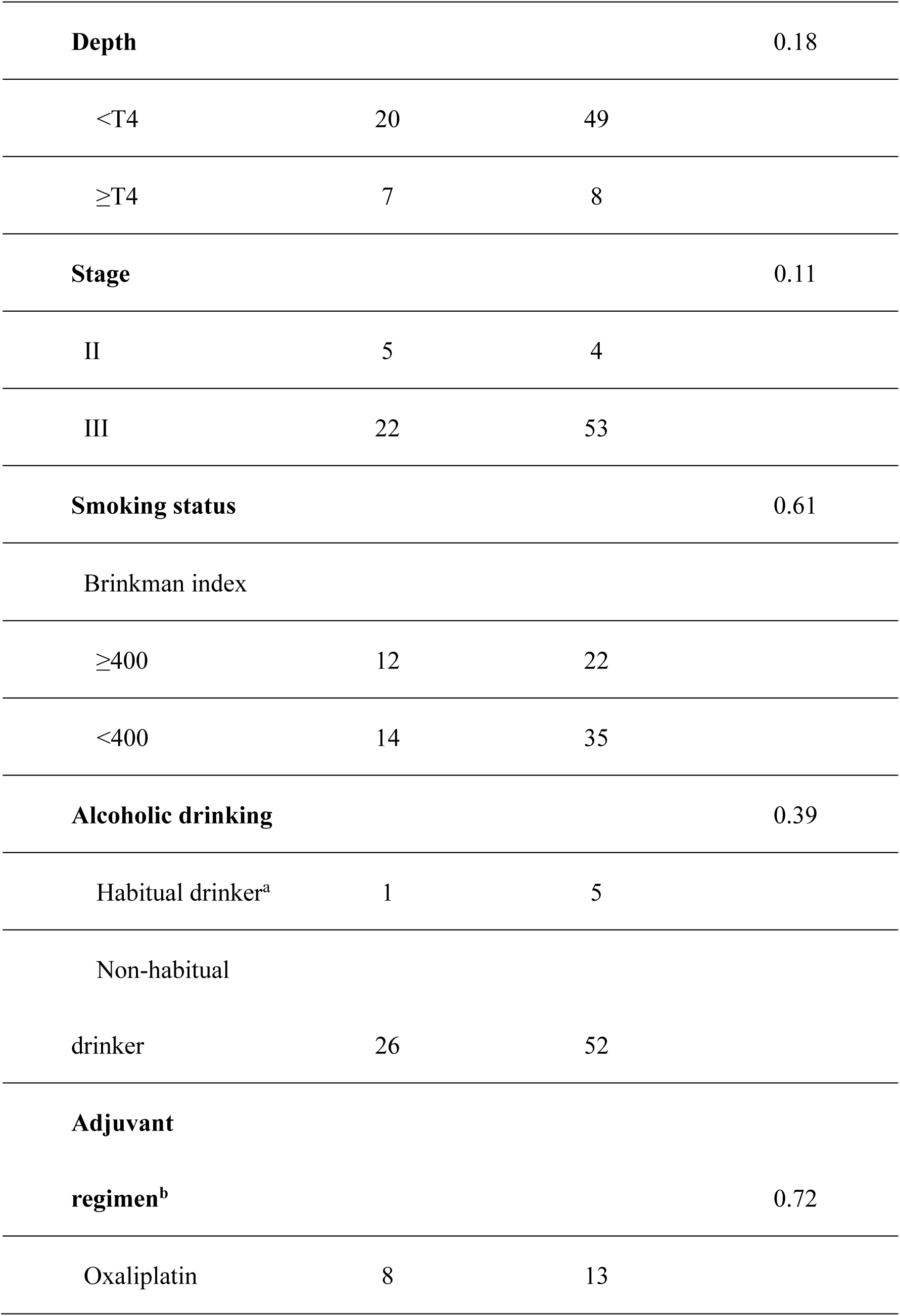
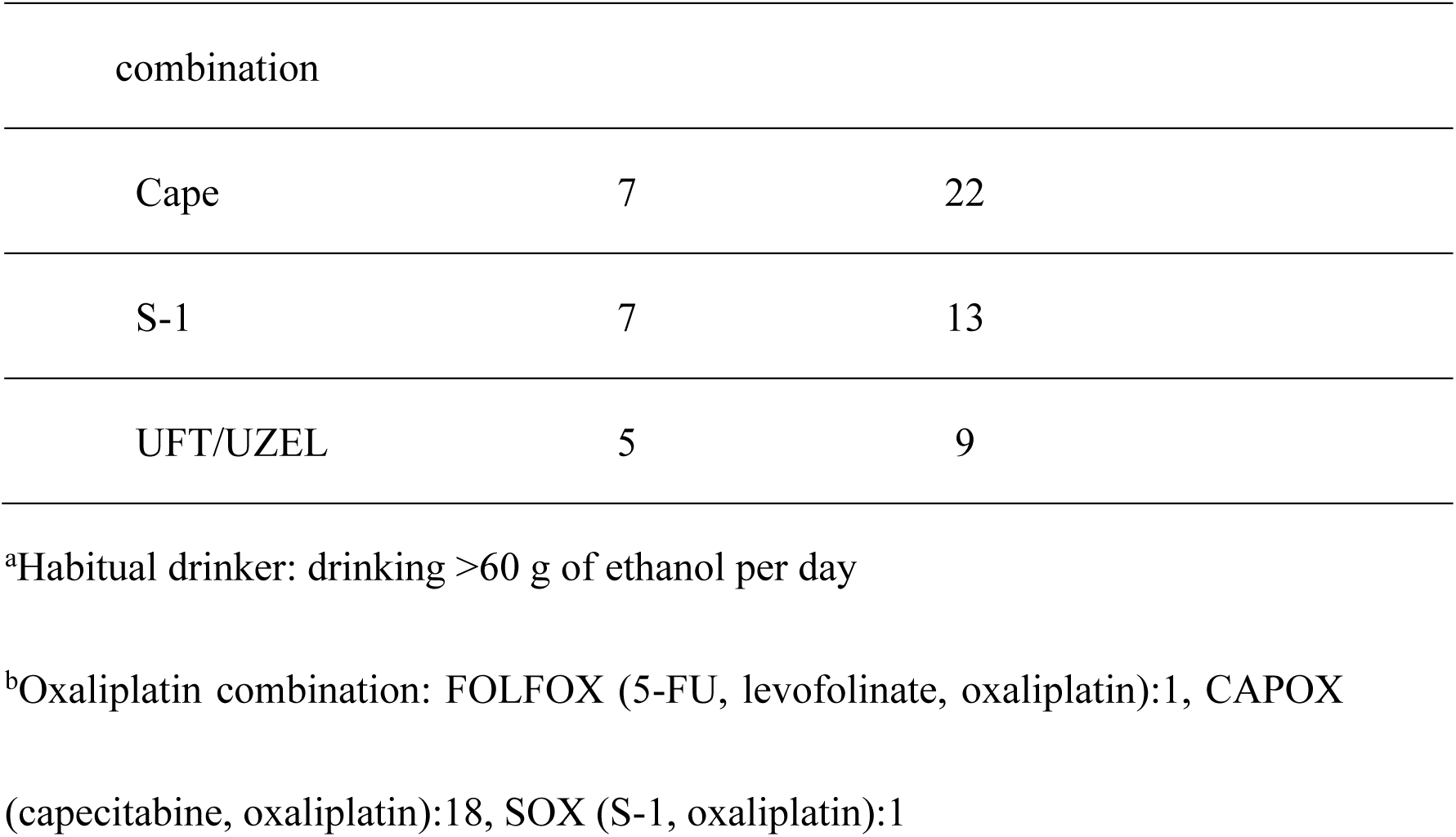
Clinical characteristics of the patients

|  | Recurrence (n = 27) | Non-recurrence (n = 57) | *p*-value |
| --- | --- | --- | --- |
| **Sex** |  |  | 0.57 |
| Male | 16 | 30 |  |
| Female | 11 | 27 |  |
| **Age (year)** |  |  | 0.8 |
| Median (range) | 65 (38–79) | 67 (40–80) |  |
| **Pathological histotype** |  |  | 0.44 |
| Non poor (tub1, tub2, pap) | 18 | 33 |  |
| Poor (por, muc) | 9 | 24 |  |
| **Location** |  |  | 0.49 |
| Right | 7 | 19 |  |
| Left | 20 | 38 |  |
| **Depth** |  |  | 0.18 |
| <T4 | 20 | 49 |  |
| ≥T4 | 7 | 8 |  |
| **Stage** |  |  | 0.11 |
| II | 5 | 4 |  |
| III | 22 | 53 |  |
| **Smoking status** |  |  | 0.61 |
| Brinkman index |  |  |  |
| ≥400 | 12 | 22 |  |
| <400 | 14 | 35 |  |
| **Alcoholic drinking** |  |  | 0.39 |
| Habitual drinker<sup>a</sup> | 1 | 5 |  |
| Non-habitual |  |  |  |
| drinker | 26 | 52 |  |
| **Adjuvant** |  |  |  |
| **regimen<sup>b</sup>** |  |  | 0.72 |
| Oxaliplatin | 8 | 13 |  |

| combination |  |  |
| --- | --- | --- |
| Cape | 7 | 22 |
| S-1 | 7 | 13 |
| UFT/UZEL | 5 | 9 |

<sup>a</sup>Habitual drinker: drinking >60 g of ethanol per day
<sup>b</sup>Oxaliplatin combination: FOLFOX (5-FU, levofolinate, oxaliplatin):1, CAPOX
(capecitabine, oxaliplatin):18, SOX (S-1, oxaliplatin):1

### Statistical analysis

Fisher’s exact test was performed to determine significant associations between gene SNVs and cancer recurrence and non-recurrence (R package; https://bioconductor.org/packages/release/-bioc/html/edgeR.html). Logistic regression was performed to validate confounding factors using JMP Pro 16 (SAS Institute Inc., Cary, NC, USA). All statistical tests were two-sided, and *p* < 0.05 was considered significant.

### DNA extraction, quantification, and quality control

Samples from 84 patients were analyzed. The assessment and recovery of the cancerous areas were performed using previously reported methods.^19^ Chromosomal DNA was isolated from formalin-fixed paraffin embedded (FFPE) colorectal adenocarcinoma samples using the QIAamp DNA FFPE Tissue Kit (QIAGEN, Hilden, Germany) according to the manufacturer’s instructions. DNA concentration was determined by fluorescence measurements via the Qubit dsDNA kit (Thermo Fisher Scientific, Waltham, MA, USA).

### Targeted capture and sequencing

A library of the entire genomic sequence of all 50 known genes (electronic supplementary Table S1) was prepared using HaloPlex Target Enrichment kits (Agilent Technologies, Santa Clara, CA, USA) according to the manufacturer’s instructions. For each library, the confirmation of enrichment and the brief quantification of the enriched target DNA were performed using the High Sensitivity D1000 Screen Tape System (Agilent Technologies). The pooled samples with different indices for multiplex sequencing were measured using the library quantification kit (Kapa Biosystems, Wilmington, MA, USA) to obtain molar concentrations. High-throughput sequencing was performed with 150-bp paired-end reads on a MiSeq or NextSeq platform (Illumina, San Diego, CA, USA) for each pooled sample according to the manufacturers’ protocols.

### Data analysis for next generation sequencing

The raw sequence read data passed the quality checks in FastQC (http://www.bioinformatics.babraham.ac.uk/projects/fastqc). Read trimming via base quality was performed using FASTX-toolkit v.0.0.14.^20^ Read alignments to the UCSC hg38 reference genome were performed using the Burrows–Wheeler Aligner.^21^ Non-mappable reads were removed using SAMtools.^22^ After filtering these reads, the Genome Analysis Toolkit (GATK) was used for local realignment and base quality score recalibration. For SNVs and small insertion/deletion (INDEL) discovery, we applied the GATK multiple-sample calling protocol.^23^ The coverage of the targeted regions was estimated using the GATK DepthOfCoverage. In this experiment, we used SelectVariants to select variants with “DP > 10” (depth of coverage greater than 10×). The detected variants were annotated using ANNOVAR, and pathogenicity was assessed using the ClinVar_20210501 database.^24^

### Sanger sequencing analysis

The SNVs of *PINK1* (c.1018G>A and c.1562A>C) were amplified using specific primer sets. The primer set for c.1018G>A were forward primer (5′- TCGATGTGTGGTAGCCAGAG-3′) and reverse primer (5′- GATGCCCTGTTGAACCAGAT-3′). The primer set for c.1562A>C were forward primer (5′-CCGCAAATGTGCTTCATCTA-3′) and reverse primer (5′- AGCGTTTCACACTCCAGGTT-3′). PCR was performed using the PrimeSTAR Max DNA polymerase system (Takara Bio Inc., Shiga, Japan), with annealing at 53 °C (c.1018G>A) and 50 °C (c.1562A>C). Thereafter, PCR-amplified products were extracted using the QIAquick Gel Extraction Kit (QIAGEN). Sequencing in the reverse direction was undertaken according to the manufacturer’s instructions (BigDye; Applied Biosystems, Warrington, UK). Electrophoresis of the products was performed using the ABI 3500 automated DNA sequencer (Applied Biosystems).

### Online resources

The analysis and expression of the target sequences in the recurrence and non-recurrence groups are described in electronic supplementary Table S1.

## RESULTS

### Clinical characteristics

The clinical characteristics of the 84 CRC patients with adjuvant chemotherapy are shown in Table 1. Stage III patients exhibited a higher recurrence rate than Stage II patients did; however, the difference was not significant. There were no significant differences between the recurrence and non-recurrence groups. The total number of recurrence cases was 27/84 (32.1%).

### Target sequencing in our clinical CRC cases

We selected a total of 50 autophagy-related genes and CRC-associated genes and performed the identification analysis of SNVs and INDELs using targeted enrichment sequencing (electronic supplementary Table S1).

When cells are subjected to starvation conditions (including pseudo-starvation, such as in 5-FU treatment), vesicles called sequestration membranes appear in the cytoplasm. The membrane then elongates while encompassing the cytoplasm, and the tips fuse together to form autophagosomes. When the autophagosome fuses with the lysosome, the endoplasmic reticulum is degraded, and the amino acids obtained by self-digestion are recycled as a nutrient source.^15, 25^ Several genes are required for the formation of autophagosomes, and they can be broadly classified into the following functional groups: three genes (*PIK3R4*, *BECN1*, and *ATG14*) that contribute to the “Vps34 PI3 kinase complex” formation, seven genes (*MAP1LC3A*, *ATG3*, *ATG4A*, *ATG4B*, *ATG4C*, *ATG4D*, and *ATG7*) that contribute to the “Atg8-conjugation system,” five genes (*ATG5*, *ATG10*, *ATG12*, *ATG16L1*, and *ATG16L2*) involved in the “Atg12- conjugation system,” eight genes (*ULK1*, *ATG13*, *RB1CC1*, *MTOR*, *RPTOR*, *DEPTOR*, *AKT1S1*, and *PTEN*) that are needed for the formation of the “Atg1 protein kinase complex,” and two genes (*ATG9A* and *ATG9B*) that are important for the “Atg9 and Atg2-Atg18 complex.”^25^ In addition, mitophagy is a selective degradation mechanism of mitochondria via autophagy and is involved in the metabolism of old mitochondria. Eight genes (*PINK1*, *PRKN*, *BNIP3*, *BNIP3L*, *FUNDC1*, *OPTN*, *BCL2L13*, and *CALCOCO2*) contribute to the “mitophagy receptor.”^25^ It was reported that *KRAS*- induced autophagy proceeds via the upregulation of the MEK/ERK pathway in colon models and that *KRAS* and autophagy contribute to CRC cell survival during starvation. Ten genes (*KRAS*, *NRAS*, *HRAS*, *ARAF*, *BRAF*, *RAF1*, *MAP2K1*, *MAP2K2*, *MAPK1*, and *MAPK3*) contribute to the “RAS-MEK/ERK pathway.”^26^ Six genes (*APC*, *CTNNB1*, *ERBB2*, *SMAD4*, *PIK3CA*, and *TP53*) recognized to be mutated in CRC were selected as CRC-related genes.^27^

Target regions were designed to enrich the exonic regions and exon–intron junctions of all 50 genes (electronic supplementary Table S1). The mean percentile of covered target regions was 98.49%.

### Quality assessment

A median of 3,493,323 sequence-mapped reads were obtained per sample (range: 437,058–6,034,492 reads/sample). Among the designed target bases, 93.2% (range: 58.3%–97.9% per sample) had at least a 10-fold coverage, with a mean coverage of 1,003-fold (range: 71- to 3,342-fold) per nucleotide in the coding region of the target gene (Figures 1a and 1b). Although one sample with a significantly low coverage was found, it was not expected to have a significant effect on the overall results because the GATK multiple-sample calling protocol was applied to SNVs and small INDEL discovery in the sequencing analysis. Furthermore, the SelectVariants option was applied to remove data with a depth of coverage less than 10.

**FIG. 1.**
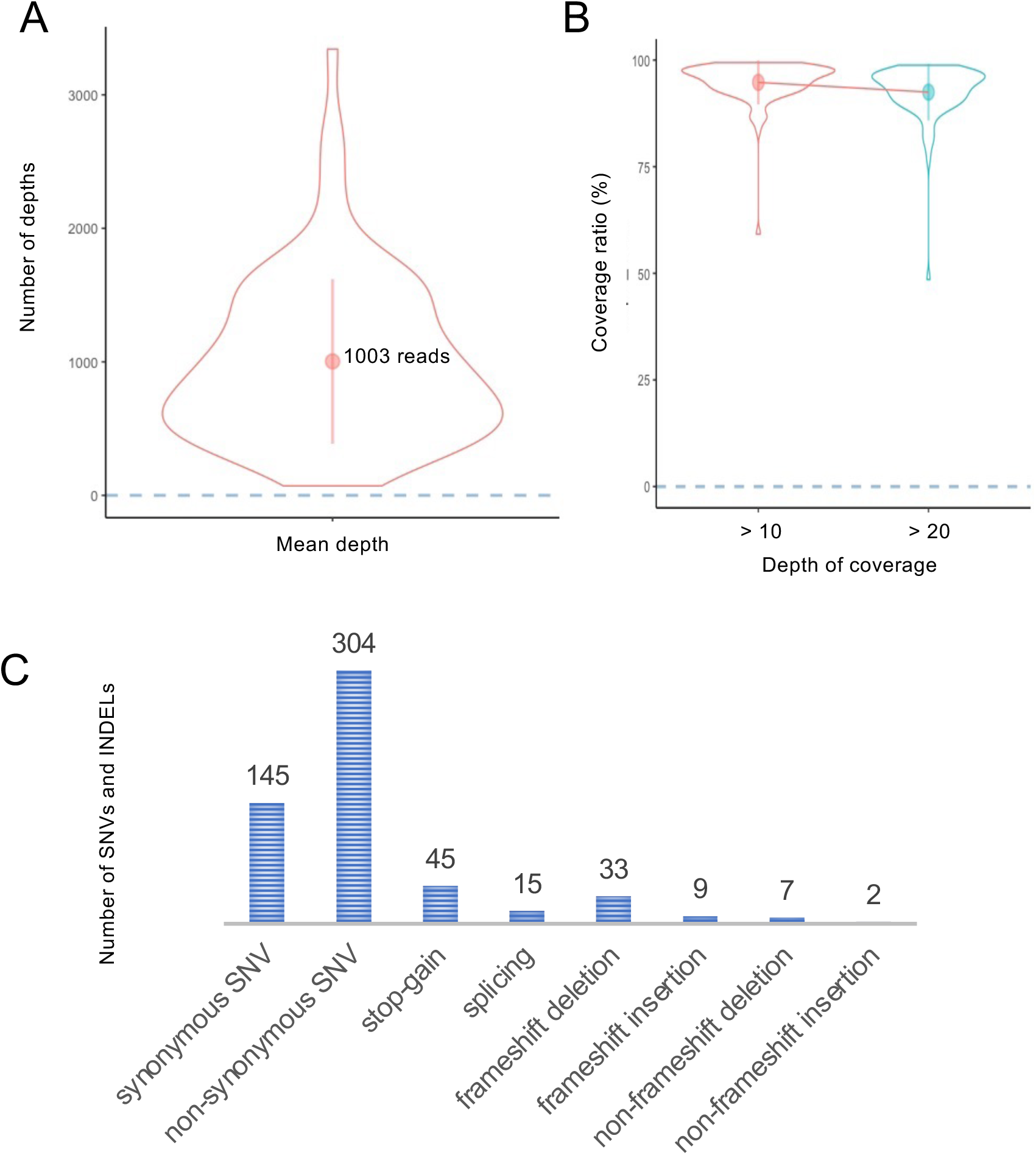
Results of the original target enrichment sequencing in the clinical cases of older adults. (a) The trend of the mean depth for each of the 84 multiplexed samples is shown in the violin plot. (b) The trend of the coverage ratio for each of the 84 multiplexed samples is shown in the violin plot. (c) The number of SNVs or INDELs identified by the original target enrichment sequencing is shown. Classification was performed by variant type. SNV: single-nucleotide variant, INDEL: insertion/deletion

### Breakdown of SNVs and INDELs

The original target enrichment sequencing for cases with postoperative adjuvant chemotherapy showed that 560 SNVs or INDELs were detected in the target region (Figure 1c). There were 304 non-synonymous SNVs in the amino acid sequences: 33 had frameshift deletions and 9 had frameshift insertions (Figure 1c). SNVs showing the stop-gain variant were found in 45 locations (Figure 1c).

### Results for individual SNVs obtained by target enrichment sequencing

The samples of 84 patients were included in the analysis. Fisher’s exact test performed on the 560 SNVs or INDELs indicated that the variants were lower in the recurrence group (n = 27) than in the non-recurrence group (n = 57). A significant difference of less than *p* < 0.05 was found in two non-synonymous SNVs: *PINK1* c.1018G>A and *PINK1* c.1562A>C and three synonymous SNVs (*KRAS* c.519T>C, *DEPTOR* c.135C>T, and OPTN c.102G>A) (Table 2). However, neither the c.1018G>A nor the c.1562A>C SNV in *PINK1* showed a statistically significant relationship with OS (Figures 2a and 2b). Logistic regression was performed to test for confounding effects on the relationship between the two *PINK1* SNVs (c.1018G>A and c.1562A>C) and the non-recurrence group, and no significant effects were found for either SNV (Tables 3 and 4).

**TABLE 2.**
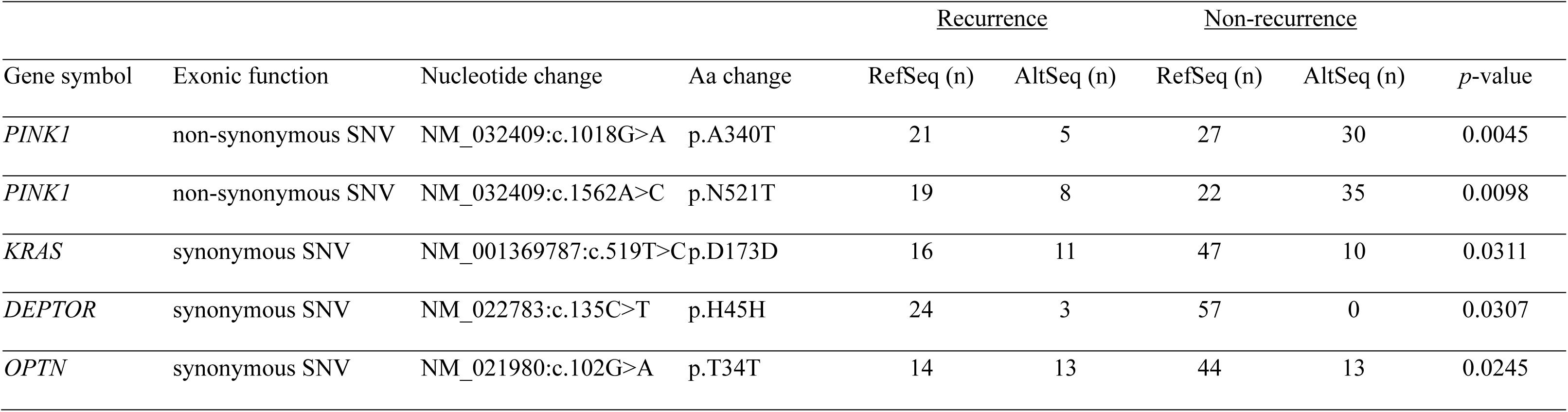
Results of target enrichment sequencing

**FIG. 2.**
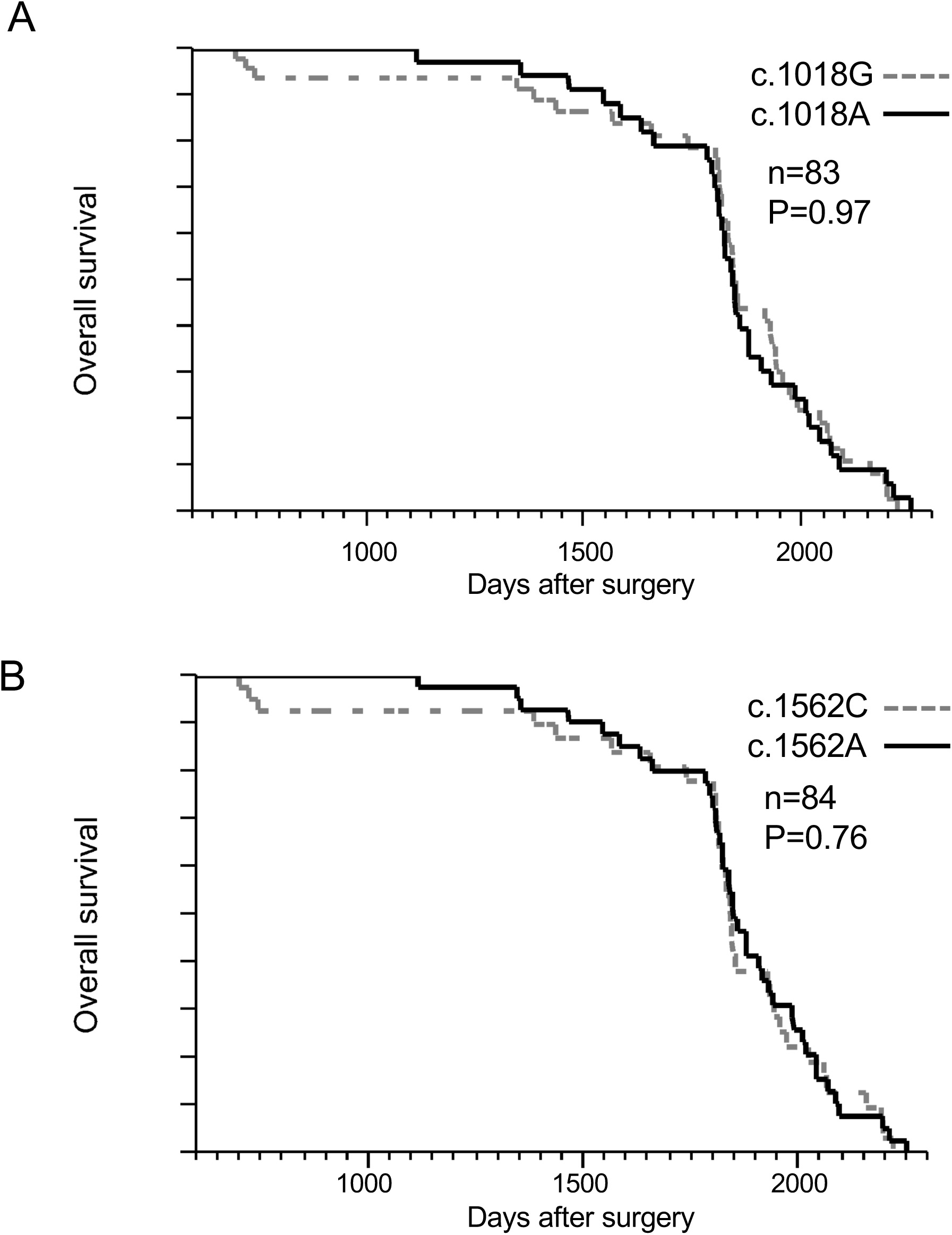
Relationship between the SNVs of *PINK1* (c.1018G>A and c.1562A>C) and CRC prognosis with 5-FU-based adjuvant chemotherapy: OS with or without c.1018G>A or (b) c.1562A>C. The analyzed specimens were 84 and 83 for (a) c.1018G>A and (b) c.1562A>C, respectively. In one case, the sample of (b) c.1018G>A was not analyzed because of an inappropriate specimen status. No statistically significant relationship was found between the two SNVs in *PINK1* and OS. CRC: colorectal cancer

**TABLE 3.**
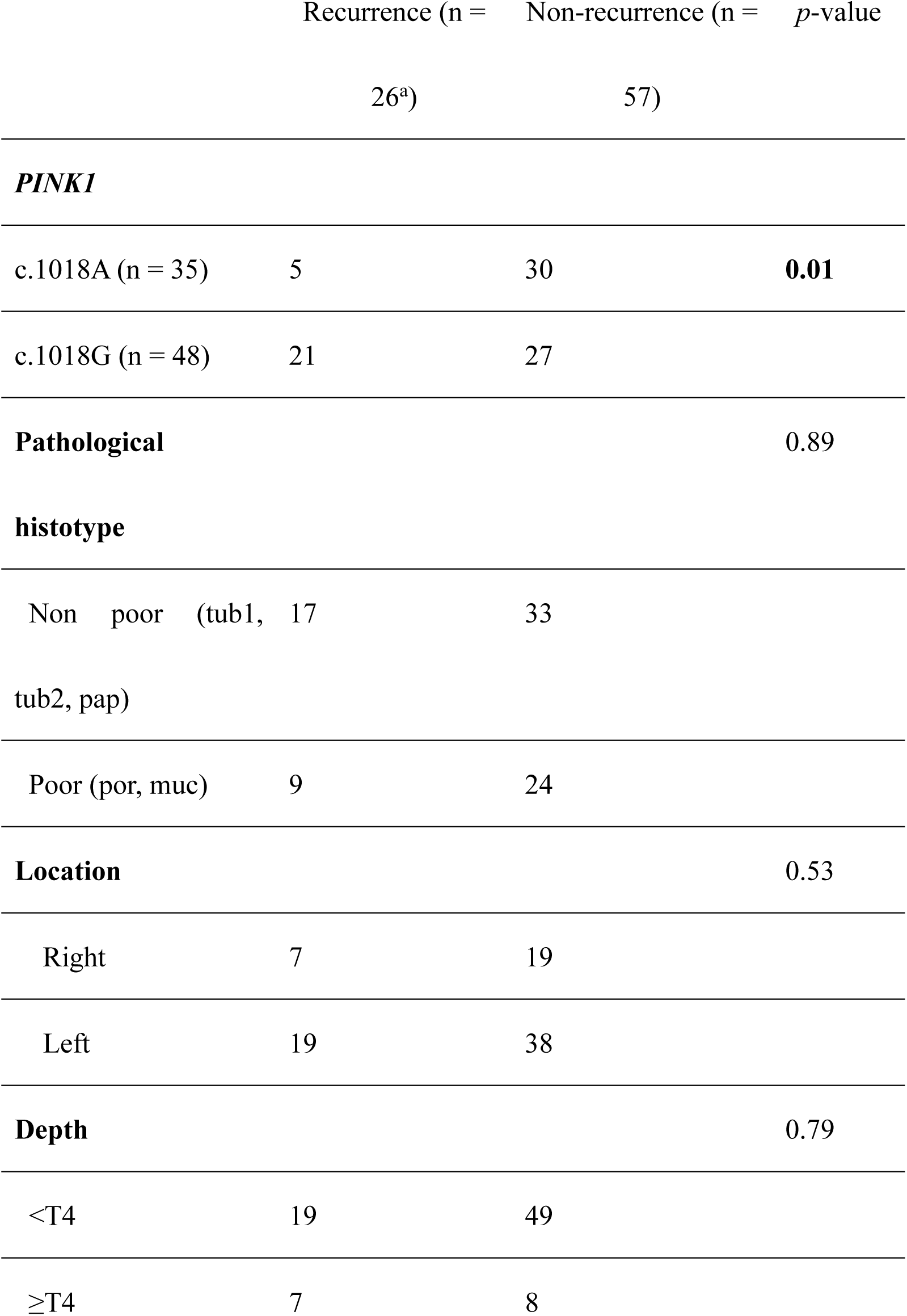
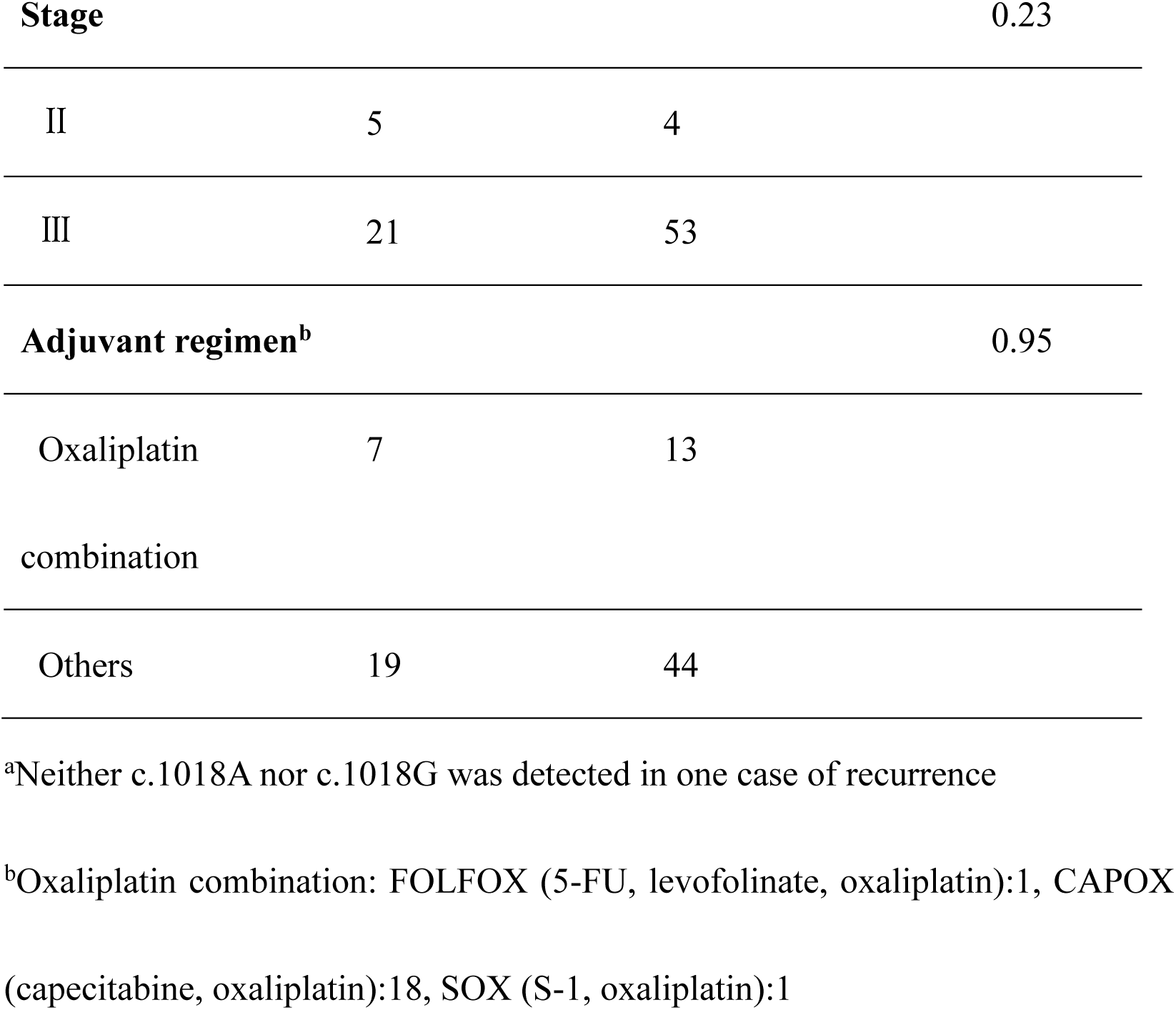
Confounding factors influencing the recurrence prevention effectiveness of an SNV in *PINK1* (c.1018G>A)

**TABLE 4.**
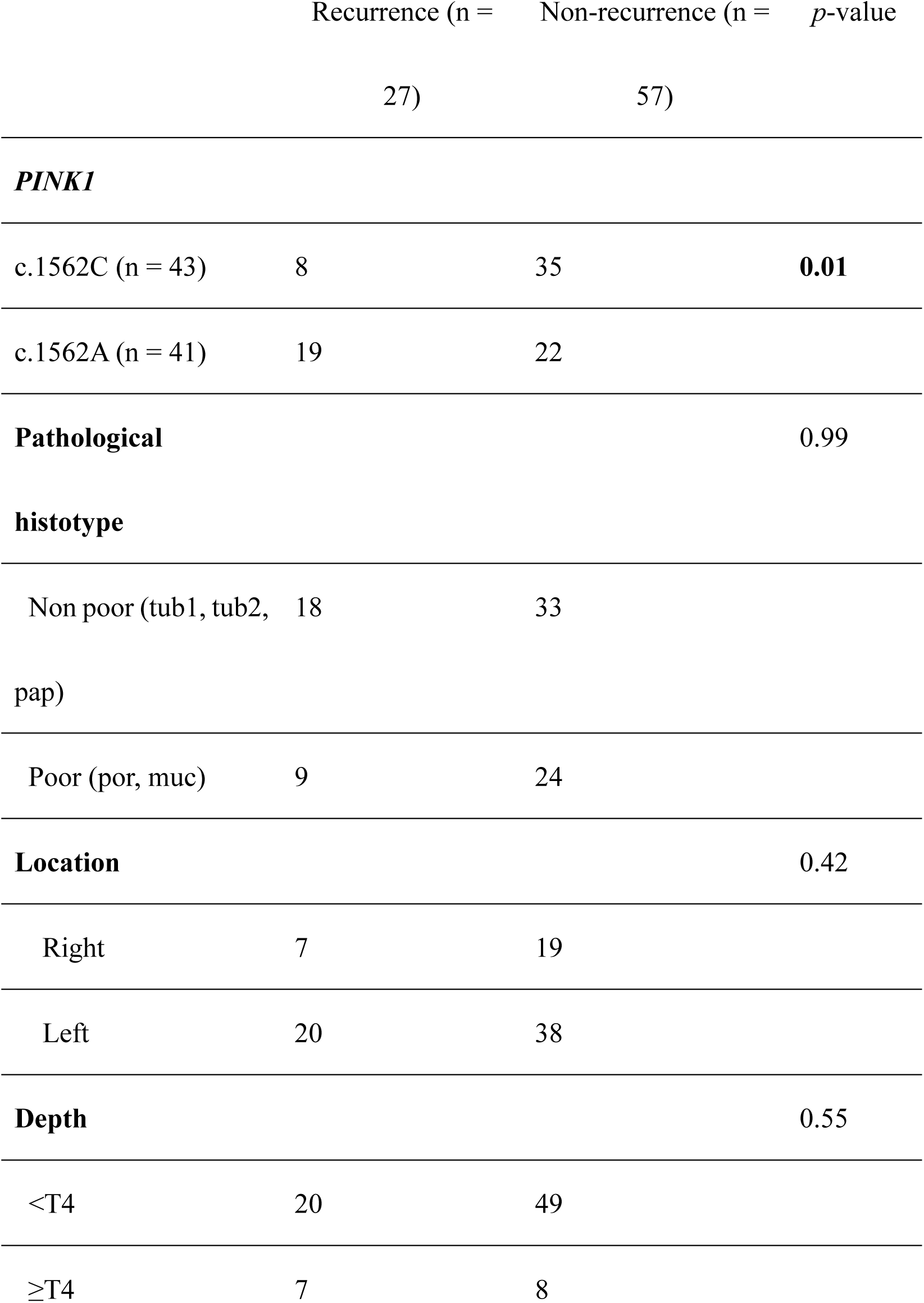
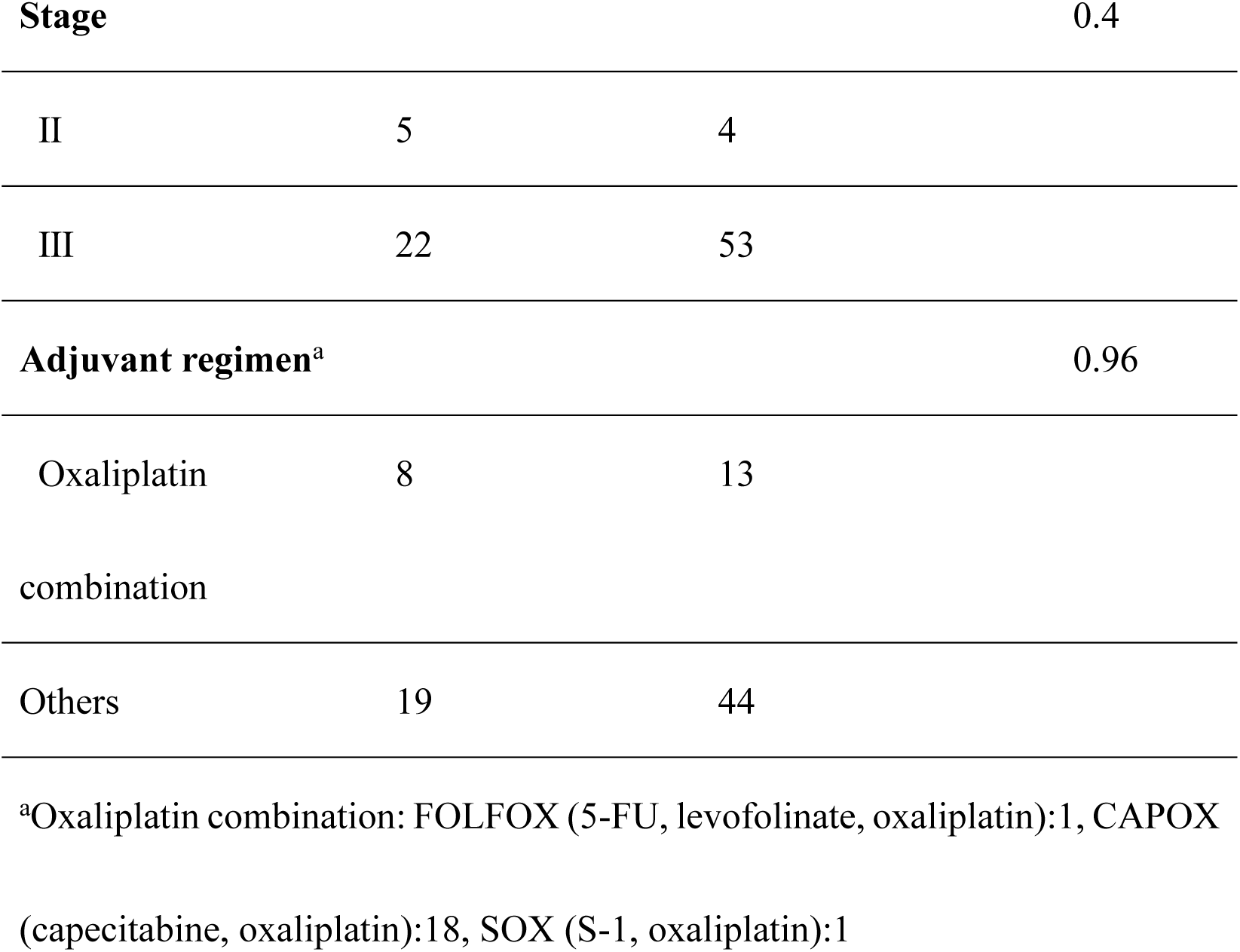
Confounding factors influencing the recurrence prevention effectiveness of an SNV in PINK1 (c.1562A>C)

### SNVs of PINK1 gene

The PINK1/Parkin pathway is the most studied pathway of mitophagy, and the serine/threonine PINK1 is the initiator of this pathway.^28^ In this target enrichment sequencing, five non-synonymous SNVs of PINK1 were found in the kinase domains (KDs) (Figure 3a). One of the remaining SNVs was also found on the C-terminal domain (CTD) sequence (Figure 3a), which controls the structure of the KD and helps the kinase region identify the substrate.^29^ In the current study, two SNVs showed significant differences, but we considered that significant differences might have been observed in other SNVs on the KD if the number of cases had been higher.

**FIG. 3.**
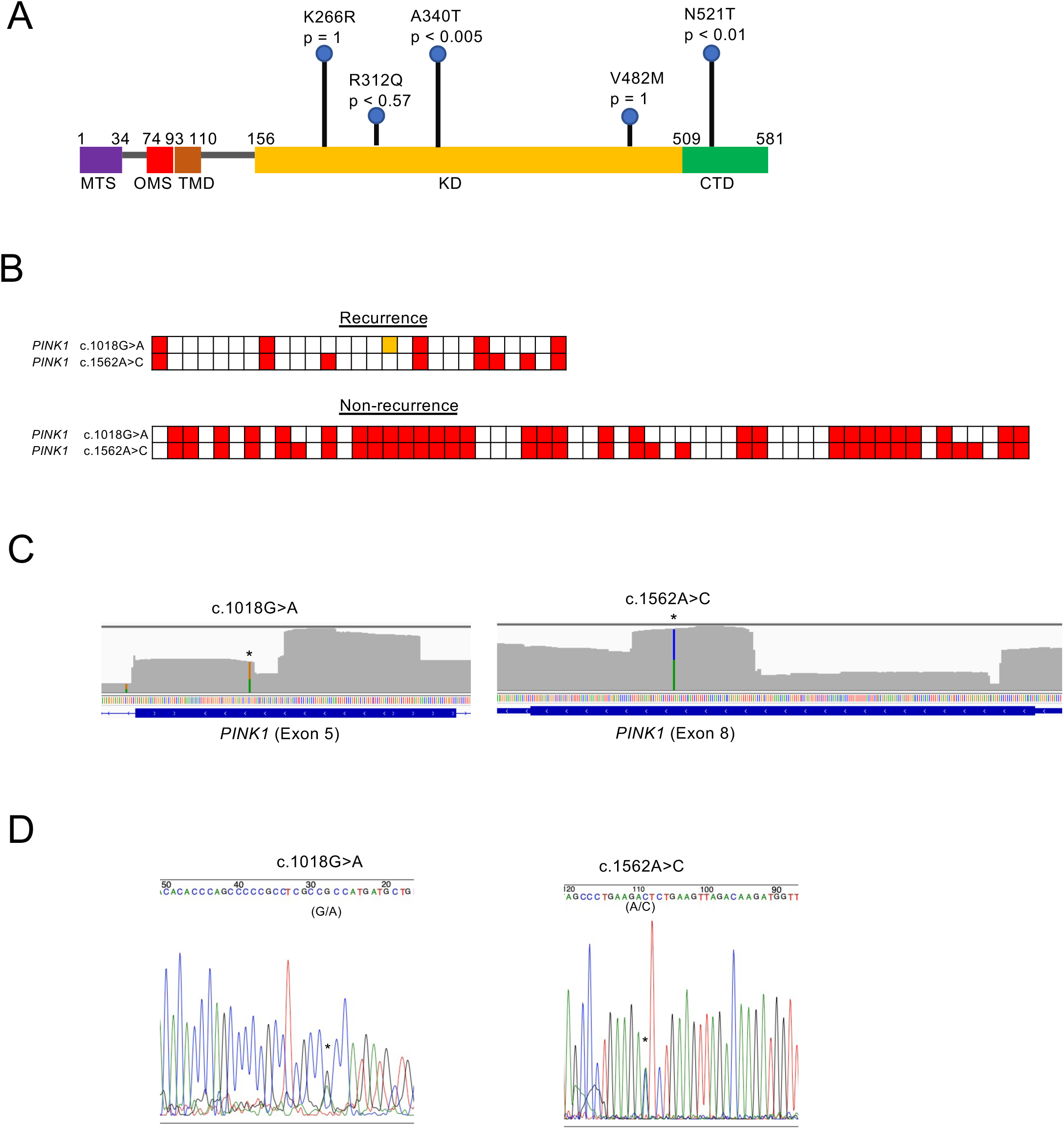
SNVs of *PINK1* gene detected in this study. (a) The full-length PINK1 can be divided into five structural and functional regions: MTS, OMS, TMD, KD, and CTD. p indicates the *p*-value. CTD, C-terminal domain; KD, kinase domain; MTS, mitochondrial targeting sequence; OMS, outer membrane localization signal; TMD, transmembrane domain. (b) Samples with AltSeq for *PINK1* are shown in red. Samples whose sequence reads did not satisfy the criteria are marked in yellow. (c-d) Sequencing chromatograms illustrating PINK1 c.1018G>A and c.1562A>C. * indicates the location of the SNV. (c) Integrative Genomics Viewer screenshot of soft-clipped reads in exons 5 and 8 of PINK1. Green represents A, yellow represents G, and blue represents C. (d) Sanger sequencing data.

*PINK1* was found in two SNVs (c.1018G>A and c.1562A>C) with significance in the non-recurrence group. The SNVs were marked in red for each patient in which they were found, and c.1018G>A was always found in conjunction with c.1562A>C (Figure 3b).

The results obtained by the next generation sequencing of c.1562A>C and c.1018G>A of *PINK1* were displayed using the Integrative Genomics Viewer software (Figure 3c). The same positions were reconfirmed by the Sanger sequencing method, and the overlap of G (black) and A (green) waveforms was observed in c.1018, as well as in A (green) and C (blue) waveforms in c.1562 (Figure 3d).

### Correlation between pathogenic/likely pathogenic SNV and recurrence rate

Fisher’s exact test was performed in the recurrence group and the non-recurrence group for each gene that showed non-synonymous SNVs resulting in amino acid substitutions or SNVs with significant effects such as frameshift deletion, frameshift insertion, stop-gain, and splicing. The results showed significant differences only for *PINK1* (electronic supplementary Table S2, Figure 3a).

Fisher’s exact test was performed for each gene that showed non-synonymous SNVs and was determined to be pathogenic/likely pathogenic in ClinVar, and no genes were significantly different (electronic supplementary Table S3).

## DISCUSSION

The c.1018G>A and c.1562A>C of the mitophagy-related gene *PINK1* may be used as biomarkers of non-recurrence in postoperative adjuvant chemotherapy for CRC. Although there is a worldwide consensus on postoperative adjuvant chemotherapy for stage III and high-risk stage II CRC, several problems still exist. First, criteria for high-risk stage II CRC vary among academic societies. Second, adjuvant chemotherapy with 5-FU has limited efficacy in preventing relapse.^2, 8^ Third, 5-FU-based adjuvant chemotherapy causes severe toxicity.^9, 10^ The current study does not provide a direct solution to these problems. However, the development of biomarkers for predicting the efficacy of adjuvant chemotherapy would avoid unnecessary administration, improve patients’ quality of life, and reduce costs, thus resulting in indirect solutions. In our search for biomarkers, we focused on mitophagy-related genes because mitophagy has recently been associated with chemotherapy resistance.

Mitophagy is a unique autophagic action that removes damaged mitochondria and plays an important role in maintaining mitochondrial quality. The mitophagy pathways can be broadly classified into the PINK1-Parkin-mediated ubiquitin pathway or the FUNDC1/BNIP3/NIX receptor–receptor-mediated pathway.^16^ During early tumorigenesis, mitophagy inhibits tumorigenesis to protect the normal cells. However, as cancer progresses, various genetic changes occur, resulting in the accumulation of impaired mitochondria, suppression of mitophagy, and promotion of tumorigenesis. Moreover, in advanced cancers, the rapid removal of mitochondria damaged by the stress of chemotherapy via mitophagy is thought to promote cancer cell survival and result in drug resistance.^17, 30^

Zhang et al. examined a total of 451 patients with unresectable colon cancer treated with FOLFIRI (5-FU, levofolinate, and irinotecan) plus bevacizumab in two phase III trials and demonstrated that some single-nucleotide polymorphisms in BNIP3 were predictors of response to the regimen.^31^ Another report analyzed 81 patients with unresectable CRC treated with 5-FU-based regimens and found that the loss of BNIP3 expression in cancer cells increased resistance to 5-FU-based drugs and worsened prognosis. These results suggested that BNIP3-related factors might predict drug resistance; however, in the current study, the SNVs of *BNIP3* were not correlated with recurrence rate. Considering that stage II colon cancer with high microsatellite instability may have a good prognosis and does not benefit from 5-FU adjuvant therapy,^32^ good prognostic factors may not necessarily be effective in preventing recurrence, and the difference between unresectable cancer and postoperative recurrence may also influence the recurrence rate. In the current study, SNVs in *PINK1* were also associated with relapse prevention but was not significantly associated with OS. No conclusions could be drawn owing to the small number of cases studied, and this study did not examine the relationship between SNVs and mutations. We believe that there was no correlation between SNVs of *KRAS* and *BRAF* and recurrence rates for the same reason.

In 159 patients with esophageal cancer who received 5-FU- and cisplatin-based preoperative chemotherapy, high expression of PINK1 was correlated with poor response to neoadjuvant chemotherapy, thus suggesting that PINK1-mediated mitophagy contributes to resistance to 5-FU-based neoadjuvant therapy.^17^

We found a correlation between several SNVs of *PINK1* (c.1018G>A and c.1562A>C) and the non-recurrence rate. If we hypothesize that the SNVs of *PINK1* reduce mitophagy activity and result in low expression, then the correlation between the SNVs of *PINK1* and lower recurrence rates is consistent with the finding that a high expression of *PINK1* in esophageal cancer during preoperative chemotherapy is correlated with poor efficacy. These results indicate that the SNVs of *PINK1* may reduce mitophagic activity, thus reducing chemotherapy resistance and enhancing the effect of 5-FU-based adjuvant. However, the biochemical significance of the two SNVs of *PINK1* has not been clarified.

This study is limited by (1) a small sample size, (2) an adjuvant chemotherapy regimen that is based on 5-FU but is not identical, (3) the lack of comparison with a surgery-alone group, and (4) incomplete functional analysis of SNVs despite them being candidate biomarkers. This is the first report of mitophagy-related SNVs as biomarkers for non-recurrence in postoperative adjuvant chemotherapy for CRC, and further research will deepen our knowledge on this topic.

In conclusion, the c.1018G>A and c.1562A>C of *PINK1* may be promising biomarkers for the favorable treatment effects of adjuvant chemotherapy in CRC. Further prospective studies are needed to understand their mechanisms of action.

## ACKNOWLEDGMENTS

We thank Drs. Shuichi Hironaka, Tsuyoshi Fukumoto, and Yosuke Mizuno, Mrs. Saori Koyano, and the staff of the Division of Analytical Science, Hidaka Branch of Biomedical Research Center, Saitama Medical University for providing research equipment and offering important advice.

## ETHICS DECLARATIONS

### Ethical Approval

This study was approved by the Institutional Review Board of SMUIMC (20-081). The requirement for informed consent was waived by the Institutional Review Board of SMUIMC (20-081) because of the retrospective nature of the study.

## List of supporting information

**Supplementary TABLE S1.**
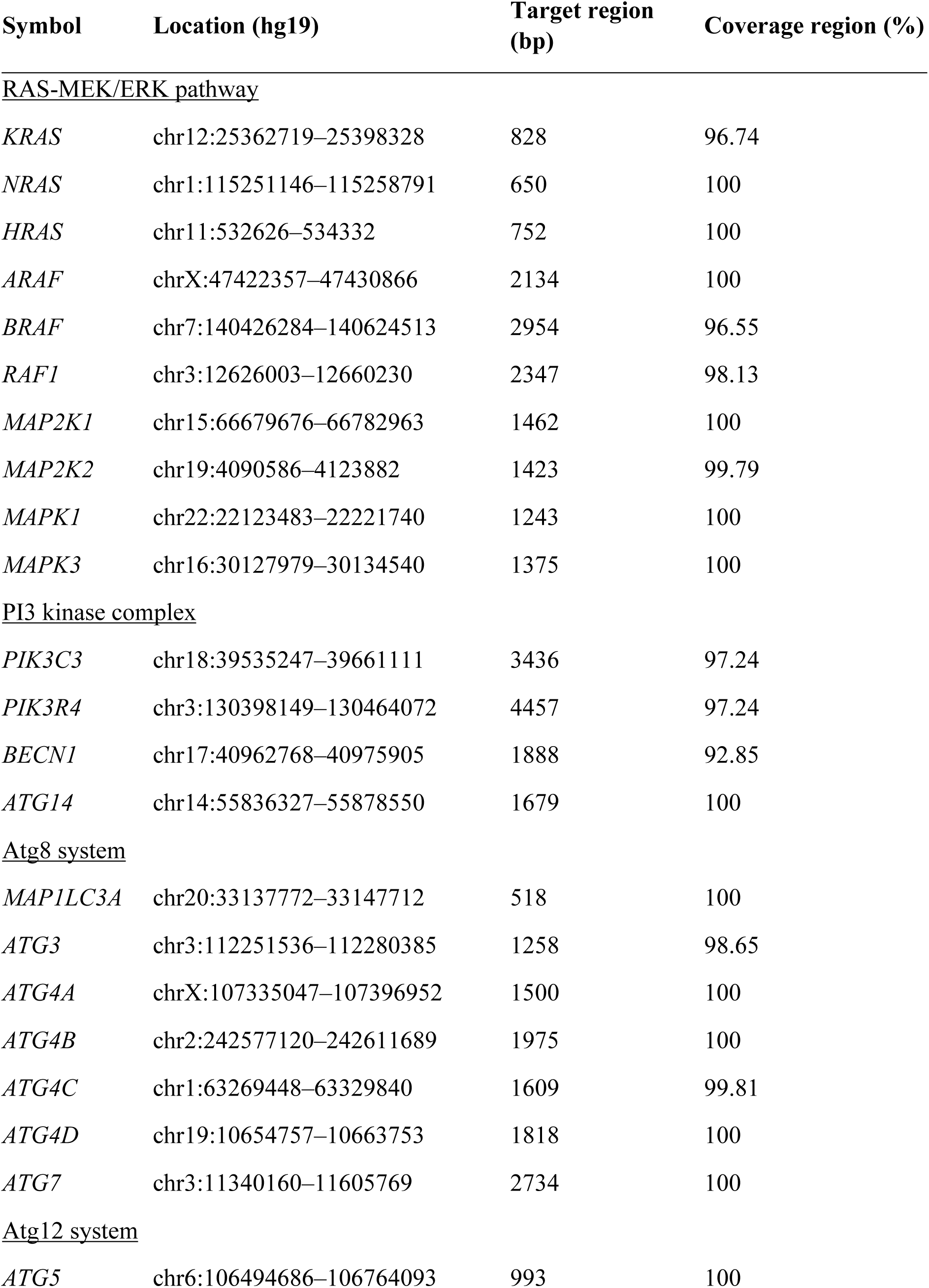
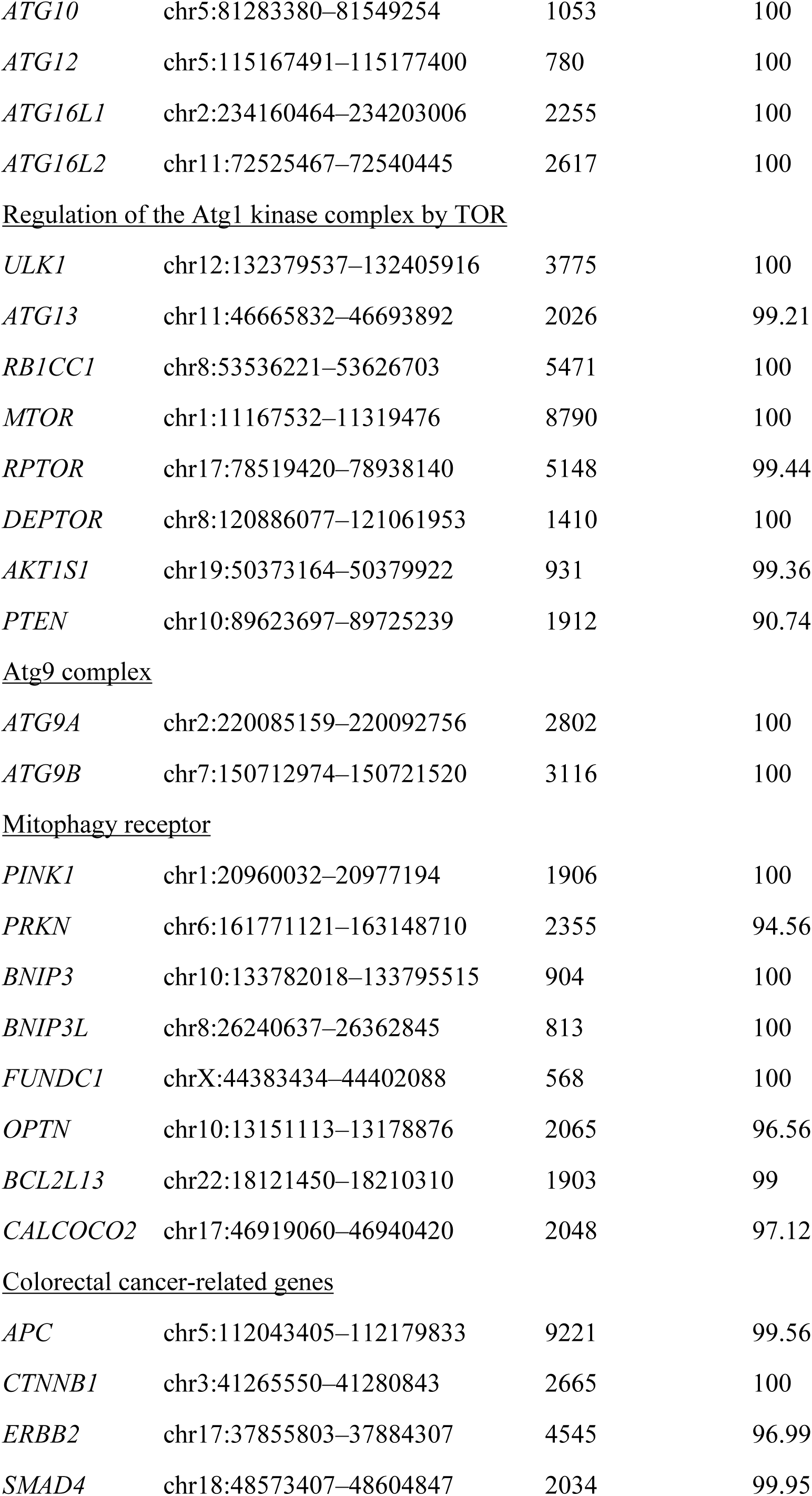
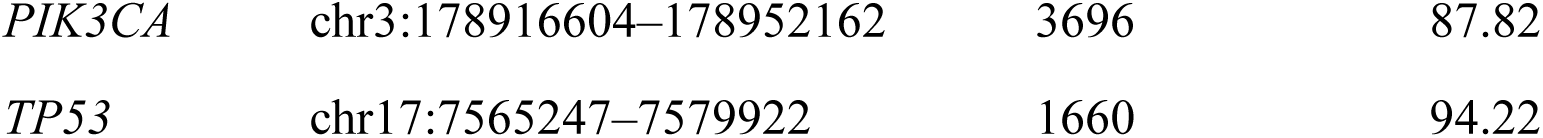
Targeted sequences in the selected clinical colorectal cancer cases

**Supplementary TABLE S2.**
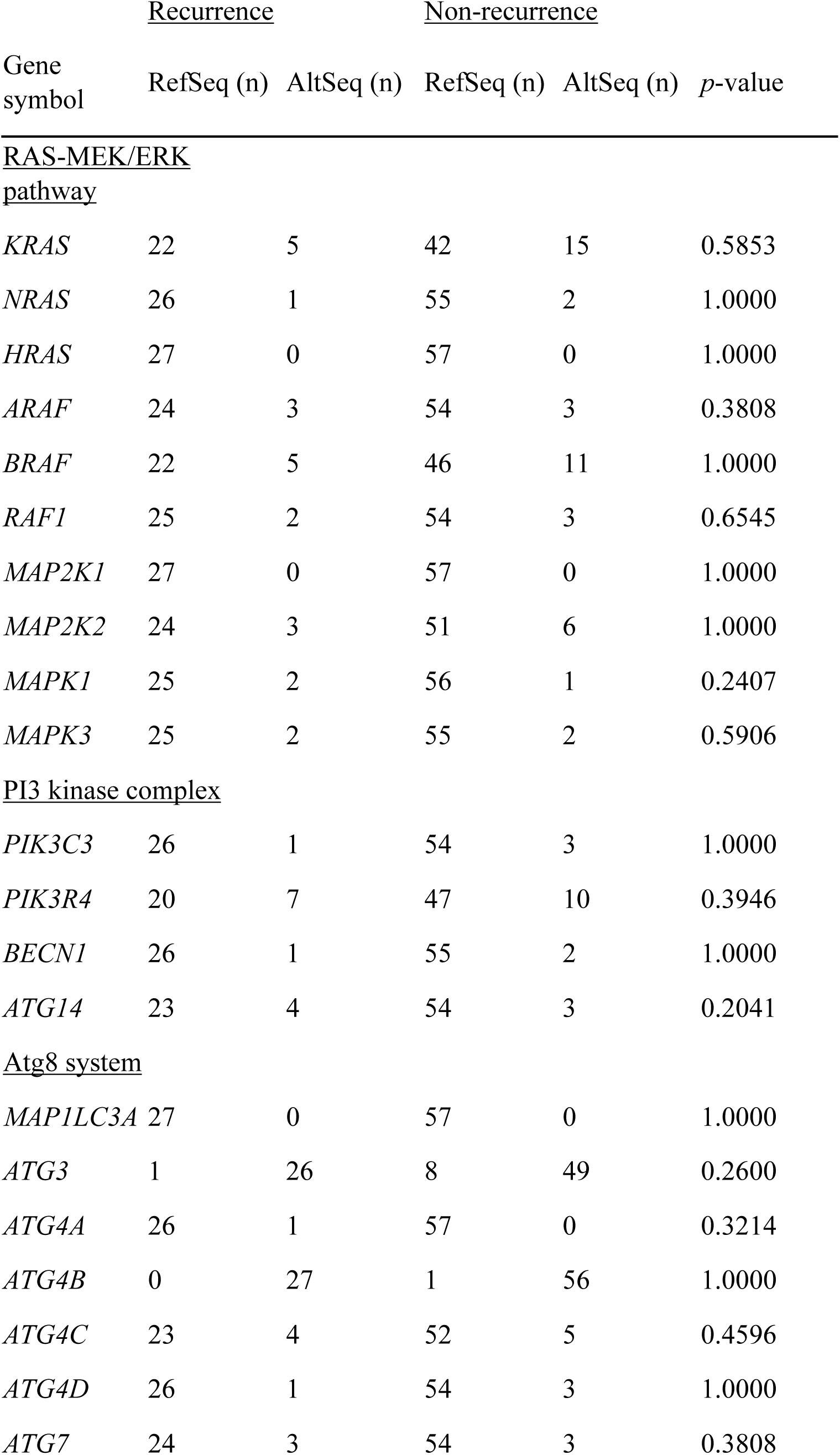
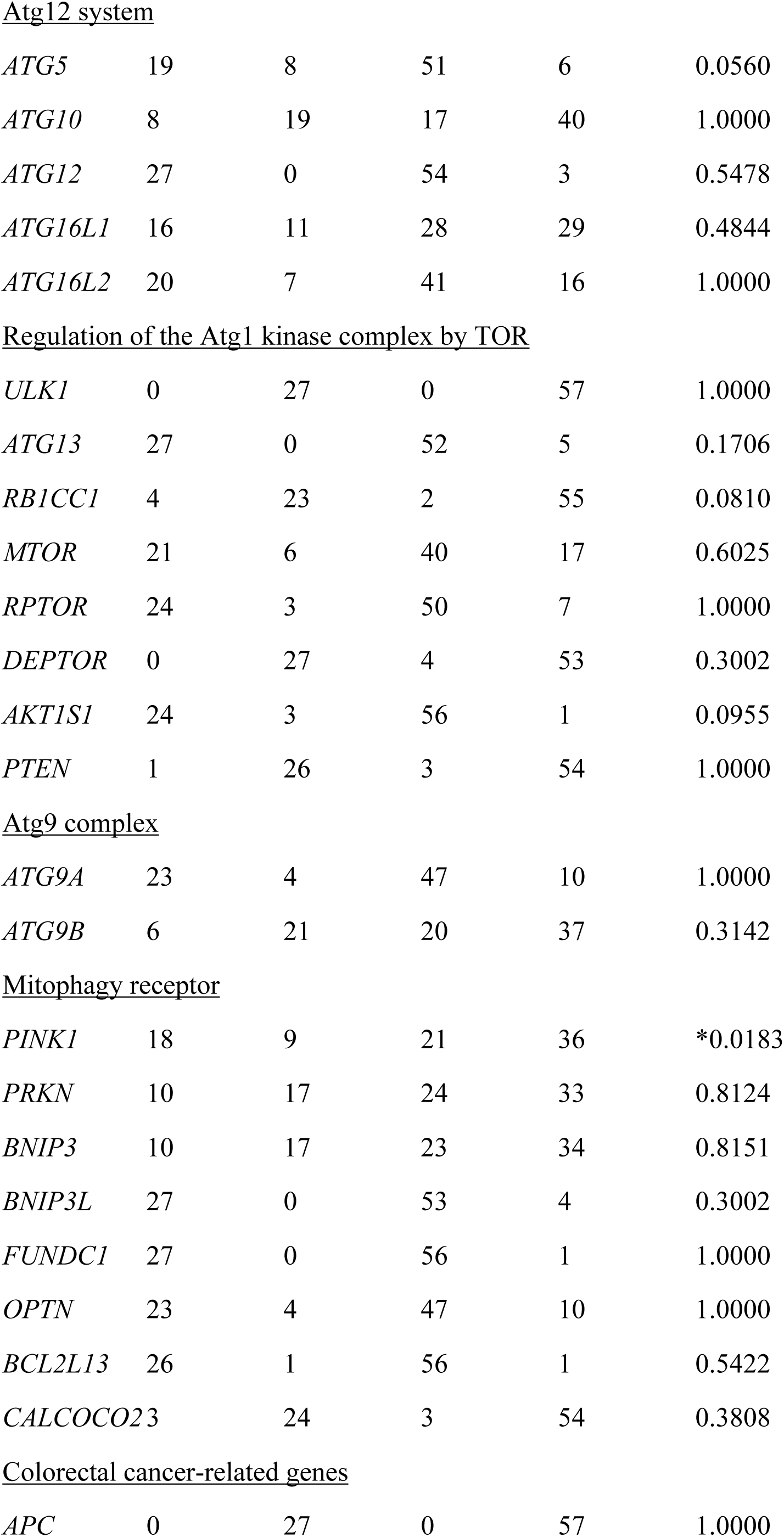
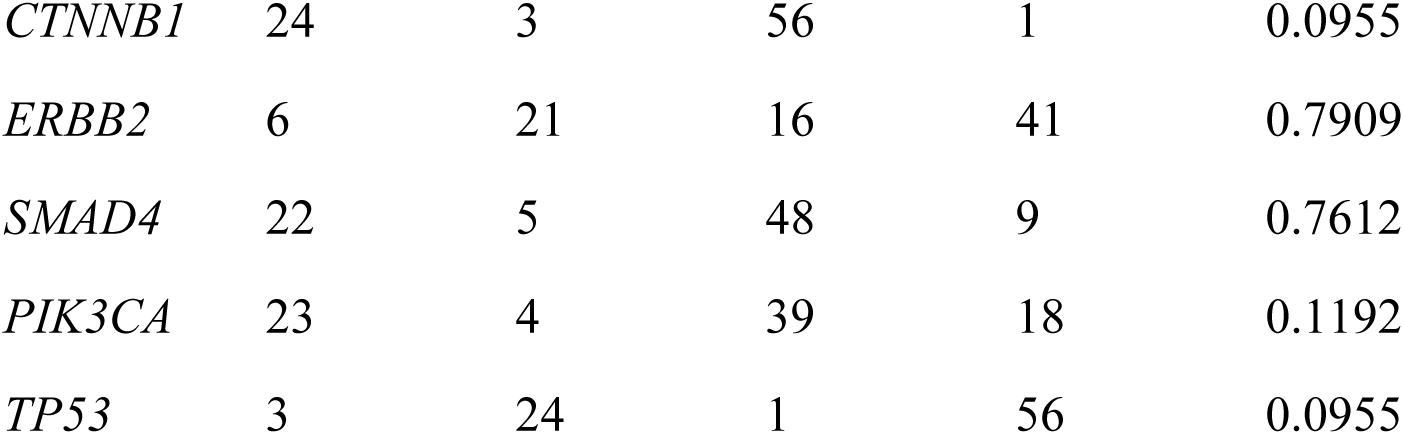
Correlation between all SNVs and recurrence rate

**Supplementary TABLE S3.** Correlation between ClinVar-based pathogenic SNVs and recurrence rate

| Gene symbol | <u>Recurrence</u> |  | <u>Non-recurrence</u> |  | *p*-value |
| --- | --- | --- | --- | --- | --- |
|  | RefSeq (n) | AltSeq (n) | RefSeq (n) | AltSeq (n) |  |
| *APC* | 8 | 19 | 19 | 38 | 0.8063 |
| *BRAF* | 26 | 1 | 53 | 4 | 1.0000 |
| *CTNNB1* | 27 | 0 | 57 | 0 | 1.0000 |
| *KRAS* | 22 | 5 | 45 | 12 | 1.0000 |
| *NRAS* | 27 | 0 | 55 | 2 | 1.0000 |
| *PIK3CA* | 25 | 2 | 45 | 12 | 0.2082 |
| *PTEN* | 27 | 0 | 56 | 1 | 1.0000 |
| *RAF1* | 27 | 0 | 56 | 1 | 1.0000 |
| *SMAD4* | 25 | 2 | 54 | 3 | 0.6545 |
| *TP53* | 10 | 17 | 30 | 27 | 0.2431 |

## REFERENCES

1. Sung H, Ferlay J, Siegel RL, et al. Global cancer statistics 2020: GLOBOCAN estimates of incidence and mortality worldwide for 36 cancers in 185 countries. CA Cancer J Clin. 2021;71:209–249.

2. O’Connell MJ, Campbell ME, Goldberg RM, et al. Survival following recurrence in stage II and III colon cancer: findings from the ACCENT data set. J Clin Oncol. 2008;26:2336–2341.

3. Sargent D, Sobrero A, Grothey A, et al. Evidence for cure by adjuvant therapy in colon cancer: observations based on individual patient data from 20,898 patients on 18 randomized trials. J Clin Oncol. 2009;27:872–877.

4. Benson AB, Schrag D, Somerfield MR, et al. American Society of Clinical Oncology recommendations on adjuvant chemotherapy for stage II colon cancer. J Clin Oncol. 2004;22:3408–3419.

5. Labianca R, Nordlinger B, Beretta GD, et al. (2013) Early colon cancer: ESMO Clinical Practice Guidelines for diagnosis, treatment and follow-up. Ann Oncol. 2013;24(Suppl 6):VI64–VI72.

6. Benson AB, Venook AP, Al-Hawary MM, et al. Colon cancer, Version 2.2021, NCCN clinical practice guidelines in oncology. J Natl Compr Canc Netw. 2021;19:329–359.

7. Tie J, Wang Y, Tomasetti C, et al. Circulating tumor DNA analysis detects minimal residual disease and predicts recurrence in patients with stage II colon cancer. Sci Transl Med 2016;8:346ra92.

8. Tie J, Cohen JD, Lahouel K, et al. Circulating tumor DNA analysis guiding adjuvant therapy in stage II colon cancer. N Engl J Med. 2022;386:2261–2272.

9. Sargent DJ, Goldberg RM, Jacobson SD, et al. A pooled analysis of adjuvant chemotherapy for resected colon cancer in elderly patients. N Engl J Med. 2001;345:1091–1097.

10. Yokoi K, Nakajima Y, Matsuoka H, et al. Impact of *DPYD*, *DPYS*, and *UPB1* gene variations on severe drug-related toxicity in patients with cancer. Cancer Sci. 2020;111:3359–3366.

11. Lunenburg CATC, Henricks LM, Guchelaar HJ, et al. Prospective *DPYD* genotyping to reduce the risk of fluoropyrimidine-induced severe toxicity: Ready for prime time. Eur J Cancer 2016;54:40–48.

12. Hashiguchi Y, Muro K, Saito Y, et al. Japanese Society for Cancer of the Colon and Rectum (JSCCR) guidelines 2019 for the treatment of colorectal cancer. Int J Clin Oncol. 2020;25:1–42.

13. Formica V, Sera F, Cremolini C, et al. *KRAS* and *BRAF* mutations in stage II and III colon cancer: a systematic review and meta-analysis. J Natl Cancer Inst. 2022;114:517–527.

14. Devenport SN, Shah YM. Functions and implications of autophagy in colon cancer. Cells 2019;8:1349.

15. Li J, Hou N, Faried A, et al. Inhibition of autophagy augments 5-fluorouracil chemotherapy in human colon cancer in vitro and in vivo model. Eur J Cancer 2010;46:1900–1909.

16. Wang Y, Liu HH, Cao YT, Zhang LL, Huang F, Yi C. The role of mitochondrial dynamics and mitophagy in carcinogenesis, metastasis and therapy. Front Cell Dev Biol. 2020;8:413.

17. Yamashita K, Miyata H, Makino T, et al. High expression of the mitophagy-related protein Pink1 is associated with a poor response to chemotherapy and a poor prognosis for patients treated with neoadjuvant chemotherapy for esophageal squamous cell carcinoma. Ann Surg Oncol. 2017;24:4025–4032.

18. He J, Pei L, Jiang H, et al. Chemoresistance of colorectal cancer to 5-fluorouracil is associated with silencing of the BNIP3 gene through aberrant methylation. J Cancer 2017;8:1187–1196.

19. Inoue H, Hirasaki M, Kogashiwa Y, et al. Predicting the radiosensitivity of HPV-negative oropharyngeal squamous cell carcinoma using miR-130b. Acta Otolaryngol. 2021;141:640–645.

20. Bolger AM, Lohse M, Usadel B. Trimmomatic: a flexible trimmer for Illumina sequence data. Bioinformatics 2014;30:2114–1120.

21. Li H, Durbin R. Fast and accurate short read alignment with Burrows–Wheeler transform. Bioinformatics 2009;25:1754–1760.

22. Li H, Handsaker B, Wysoker A, et al. The Sequence Alignment/Map format and SAMtools. Bioinformatics 2009;25:2078–2079.

23. McKenna A, Hanna M, Banks E, et al. The Genome Analysis Toolkit: a MapReduce framework for analyzing next-generation DNA sequencing data. Genome Res. 2010;20:1297–1303.

24. Wang K, Li M, Hakonarson H. ANNOVAR: functional annotation of genetic variants from high-throughput sequencing data. Nucleic Acids Res. 2010;38:e164.

25. Klionsky DJ, Abdelmohsen K, Abe A, et al. Guidelines for the use and interpretation of assays for monitoring autophagy (3rd edition). Autophagy 2016;12:1–222.

26. Alves S, Castro L, Fernandes MS, et al. Colorectal cancer-related mutant *KRAS* alleles function as positive regulators of autophagy. Oncotarget 2015;6:30787–30802.

27. Bai J, Gao J, Mao Z, et al. Genetic mutations in human rectal cancers detected by targeted sequencing. J Hum Genet. 2015;60:589–596.

28. Kulikov AV, Luchkina EA, Gogvadze V, Zhivotovsky B. Mitophagy: Link to cancer development and therapy. Biochem Biophys Res Commun. 2017;482:432–439.

29. Wang N, Zhu P, Huang R, et al. PINK1: The guard of mitochondria. Life Sci. 2020;259:118247.

30. Panigrahi DP, Praharaj PP, Bhol CS, et al. The emerging, multifaceted role of mitophagy in cancer and cancer therapeutics. Semin Cancer Biol. 2020;66:45–58.

31. Zhang S, Millstein J, Cao S, et al. Abstract 1342: Polymorphisms in genes involved in mitophagy pathway predict clinical outcome in patients (pts) with metastatic colorectal cancer (mCRC): Data from TRIBE and FIRE3 phase III trials. Cancer Res. 2019;79:1342.

32. Sargent DJ, Marsoni S, Monges G, et al. Defective mismatch repair as a predictive marker for lack of efficacy of fluorouracil-based adjuvant therapy in colon cancer. J Clin Oncol. 2010;28:3219–3226.

